# Decoding bacterial methylomes in four public health-relevant microbial species: Nanopore sequencing enables reproducible analysis of DNA modifications

**DOI:** 10.1101/2025.01.29.635492

**Authors:** Valentina Galeone, Johanna Dabernig-Heinz, Mara Lohde, Christian Brandt, Christian Kohler, Gabriel E. Wagner, Martin Hölzer

## Abstract

Investigating bacterial methylation profiles provides essential complementary information to the native DNA sequence, significantly extending our understanding of how DNA modifications influence virulence, antibiotic resistance, and the ability of bacteria to evade the immune system. Recent advancements in real-time Nanopore sequencing and basecalling algorithms have enabled the direct detection of modified bases from raw signal data, eliminating the need for bisulfite treatment of DNA. However, decoding methylation signals remains challenging due to rapid technological and methodological progress.

In this study, we focus on public health-relevant bacterial strains to analyze their methylation profiles and identify methylation motifs. Our dataset includes samples from *Staphylococcus aureus*, *Listeria monocytogenes*, *Enterococcus faecium*, and *Klebsiella pneumoniae*, sequenced on the Nanopore GridION platform using the latest flow cell chemistry (R10.4.1) and modification basecalling models (Dorado basecalling SUP model v5).

We investigated distinct methylation patterns within and between species, focusing on heavily modified genes or genomic regions. Our results reveal distinct species-specific methylation profiles, with each strain exhibiting unique modification patterns. We developed a modular pipeline using Nextflow and the Nanopore Modkit tool to streamline the detection of methylated motifs. We compared the results with outputs from MicrobeMod, a recent toolkit for exploring prokaryotic methylation and base modifications in nanopore sequencing. Our pipeline is publicly available for further use (github.com/rki-mf1/ont-methylation).

We identified known methylation motifs already described in the literature and novel *de novo* motifs, providing deeper insights into the diversity of bacterial DNA modifications. Furthermore, we identified genomic regions that are extensively methylated, which could have implications for bacterial behavior and pathogenicity. We also assess improvements in basecalling accuracy, specifically how methylated bases can influence neighboring basecalls. Recent advances in basecalling models, particularly v5 models as part of Dorado, have reduced these issues, improving the reliability of methylation detection in bacterial genomes.

In conclusion, our study highlights the potential of current nanopore sequencing tools for detecting DNA modifications in prokaryotes. By making our pipeline and results publicly available, we facilitate further research into bacterial DNA modifications and their role in microbial pathogenesis.

## Introduction

The bacterial methylome refers to the complete set of DNA methylation modifications present within a bacterial genome. These modifications primarily involve methylation of adenine at the N6 position (6mA) and cytosine at either the N4 (4mC) or C5 (5mC) positions^1^. In eukaryotic genomes, 5mC is the predominant form of DNA methylation, occurring within CpG contexts where it is tightly regulated to influence gene expression, genome stability, and chromatin structure^2^. In bacterial genomes, however, 5mC is less common and occurs alongside 6mA and 4mC, with 6mA being the most prevalent form. Furthermore, bacterial DNA methylation is motif-specific, with nearly all instances of specific motif sequences being methylated (e.g., the motif in *Escherichia coli*: 5’-G-**6mA**-TC-3’)^3,4^.

DNA methyltransferases catalyze the transfer of methyl groups from S-adenosyl-L-methionine to adenine or cytosine bases. Some bacterial DNA methyltransferases, such as Dam in *E. coli*, are conserved and active in all strains of a species, while others exhibit strain-specific variability^5^. For a long time, the most well-known and widely accepted function of these enzymes was thought to be their role in restriction-modification (RM) systems, where they protect the host genome by methylating specific sites to prevent cleavage by the cognate restriction endonuclease^6,7^. These protective systems are classified into four main groups based on their enzyme composition and mechanism of action, and these classifications also influence the characteristics of the target recognition motifs.

Type I RM systems involve a restriction endonuclease, a methyltransferase, and a specificity subunit (S-subunit). These components must assemble into a complex before they can cleave and methylate DNA. The target motifs in Type I systems are often longer and more complex with a characteristic bipartite structure, requiring the S-subunit for recognition (e.g., 5’-G**-6mA-** CNNNNNNGTC-3’). The S-subunit contains two target recognition domains (TRDs), each interacting with one half of the bipartite recognition site^8^. Type II RM systems are the most well-characterized, involving a restriction endonuclease and a separate methyltransferase that both recognize a short, palindromic sequence (e.g., 5’-G-**6mA**-TC-3’). Type III systems require a methyltransferase-restriction endonuclease complex to recognize short, non-palindromic motifs and cut outside the binding site. Finally, Type IV systems lack a methyltransferase, targeting and cleaving modified DNA^5^.

However, the discovery of so-called orphan methyltransferases challenged this traditional view^9^. These enzymes, which function independently of any restriction endonuclease, highlight that DNA methyltransferases may have additional, previously unrecognized roles in genomic regulation. Recent studies indicate that bacteria use methylation as a versatile signal in genome defense, DNA replication and repair, gene expression regulation, transposition control, and host-pathogen interactions^6,10^. In *Gammaproteobacteria* and *Alphaproteobacteria*, 6mA plays a critical role in the timing and regulation of the cell cycle, with the origins of replication exhibiting high methylation levels^11,12^. Similarly, cytosine methylation at 5′CCWGG3′ sites, catalyzed by the Dcm methyltransferase, has been shown to repress the expression of ribosomal protein genes^13^.

Recent advances in methylation detection technologies have enabled genome-wide mapping of DNA methylation in bacteria. For example, SMRT (PacBio)^14^ sequencing can directly detect DNA methylation without additional chemical treatments or amplification. Oxford Nanopore Technology (ONT) sequencing also offers methylation detection, with lower initial costs and more flexibility due to compact devices like the MinION, while yielding sequencing data for highly contiguous bacterial genome reconstruction^15,16^. In addition, ONT’s methylation detection has achieved a new level of accuracy with the introduction of the upgraded R10.4.1 flowcell^17,18^, the release of the Dorado basecaller, and improved methylation-aware basecalling models. Before Dorado, basecalling was handled by the tool Guppy, with methylation modifications inferred later using separate tools such as Megalodon or other third-party software. The introduction of Remora, also developed by ONT, marked a significant shift in this process. Unlike previous methods, Remora’s models are trained directly on raw sequencing data, and they have been integrated into Dorado, eliminating the need for separate detection steps^19^.

While nanopore sequencing accuracy and modified base detection have significantly improved^20^, methylation calling remains a challenge for the basecaller in some instances. These challenges affect not only the methylated base itself but also the surrounding bases^21^, with strain-specific errors currently limiting ONT’s applicability for high-resolution bacterial genotyping and potentially leading to the misclassification of outbreaks in diagnostic and clinical settings^22–25^. Notably, Medaka, a popular polishing tool for nanopore data^26^, has released a model designed explicitly for bacteria. Early results show that this reduces methylation-induced basecalling errors^27^.

Another consequence of ONT’s recent updates is that many tools were developed for earlier flow cell versions (e.g., R9 vs R10), and their respective models are now obsolete. While analysis pipelines for human and eukaryotic genomes are beginning to catch up, new software tools specifically developed to analyze 5mC in the context of CpG islands using the latest outputs have emerged^28–30^. However, equivalent tools and comprehensive benchmarking are still mainly lacking for bacterial genomes. One option is MicrobeMod^31^, which can annotate RM genes by cross-referencing annotations with the REBASE^32^ database and identifying methylated motifs. This gap highlights the pressing need to evaluate and develop more specialized tools and pipelines for bacterial genomes, especially in light of the impact DNA modifications can have on microbial pathogenesis^33–36^.

## Materials and methods

### Dataset

Our dataset includes isolates from four species: *Enterococcus faecium* (EF), *Klebsiella pneumoniae* (KP), *Listeria monocytogenes* (LM), and *Staphylococcus aureus* (SA). These isolates were selected from a previous study assessing the reproducibility of nanopore sequencing-based genotyping of bacterial pathogens, particularly for outbreak analysis using core genome MLST^23^.

In this first study, we initially sequenced samples with the R10.4.1 pore at 260 bp/s (4 kHz, a now discontinued translocation speed) and with Illumina technology for comparison. Sequencing and downstream analysis were performed in parallel by multiple institutes, and the results were compared to assess the reproducibility. The earlier results revealed highly strain-specific typing errors in all species, strongly associated with methylation-induced basecalling errors. To investigate these error profiles further in the original study, we selected, re-sequenced, and re-analyzed three strains per species. One strain per species was selected for good nanopore sequencing performance (matching typing results with the short-read data). Two other strains per species were selected as ‘problematic,’ exhibiting methylation-induced basecalling errors leading to high allelic differences compared to the short-read reference across all institutes. In the original study, we re-sequenced these twelve isolates by three independent institutes (LAB1, LAB2, LAB3) with R10.4.1 flow cells at the current default 400 bp/s (5 kHz) translocation speed and basecalled them with the Dorado model res_dna_r10.4.1_e8.2_400bps_sup@2023-09-22_bacterial-methylation. This allowed us to assess the impact of technical and bioinformatics advancements implemented since the initial sequencing run with 260 bp/s. In the original study, we found that nanopore data from the re-sequenced samples at 400 bp/s (5 kHz) translocation speed aligned more closely to the assemblies based on Illumina short reads, except for strains KP02 and LM46^23^. A summary of the twelve samples, including their matching to short-read data based on the original study^23^, is provided in **Supplementary Table S1**. Further details about the samples, DNA extraction, and sequencing can be found in the original study^23^.

For this study, we used and re-analyzed the nanopore raw signal data from the re-sequencing runs of the twelve samples from the three institutes generated with the current default translocation speed (400 bp/s). Please note that in this study, we also used identical sample and laboratory IDs from the original study to allow for easy comparison.

### Basecalling

We re-basecalled the raw signal data with Dorado v0.8.1 using the model dna_r10.4.1_e8.2_400bps**@**v5.0.0 with SUP (super accuracy) mode and applying the modification models 6mA and 4mC_5mC to basecall and detect modified bases.

### Assembly pipeline and annotation

We filtered reads smaller than 500 bp using Filtlong (v0.2.1)^37^. We then used Rasusa (v2.1.0)^38^ to achieve consistent 100X genome coverage for all samples; the read count of samples with higher coverage was scaled down, and samples with lower coverage were left untouched. The filtered and subsampled reads were then assembled with Flye (v2.9.5)^39^, with the “-nano-hq” option, applying the “--meta” flag in cases where it increased the size of the largest contig, representing the chromosome, or reduced the number of contigs overall. The resulting assemblies were polished using Medaka (v2.0) with the model r1041_e82_400bps_bacterial_methylation. **Table S2** provides an overview of the quality of the assemblies for each institute, including N50, number of contigs, read coverage, and whether the “--meta” flag was used. The assemblies were annotated using Bakta (v1.9.4) with default parameters^40^.

### Methylation detection using Modkit

To analyze the methylation profiles of the isolates, we aligned reads (re-basecalled using Dorado) to the corresponding genome assemblies using minimap2 (v2.28)^41^. This step included read manipulation with SAMtools (v1.21)^42^, converting the BAM files to FASTQ format while preserving methylation-related information using the parameter “-T MM, ML”. The aligned reads were also used as input for MicrobeMod (v1.0.5)^31^ for motif comparison (see method details below). We used the Modkit pileup command (v0.4.1)^43^ to aggregate information from the aligned reads into a reference-based output file that included the methylation status of all bases. Modkit was run with the optional parameter--filter-threshold set to 0.75. By default, Modkit estimates this value with the sample-probs function using the 10th percentile of confidence values within the dataset. We found that the results of this function were close to 0.75. Thus, we fixed this value as an appropriate threshold for samples and replicates to standardize our analysis across the dataset.

Methylated motifs were identified using Modkit’s motif detection function and compared with those detected by MicrobeMod. For base-level methylation analysis, Modkit provides a *Fraction Modified* value for each position, defined as: N_mod_ / N_valid_cov_, where N_valid_cov_ is the sum of reads classified as either modified or canonical (non-modified) that passed the confidence threshold. We observed that *Fraction Modified* could lead to false positives at positions with low valid coverage. To address this, we introduced a new metric *Percent Modified*, defined as: N_mod_ / (N_valid_cov_ + N_fail_ + N_diff_), where N_fail_ are the reads failing Modkit’s confidence threshold, and N_diff_ are the reads containing bases differing from the canonical base for the modification. This approach reduces false positives by counting all reads, not just those passing the confidence threshold. *Percent Modified,* as a measure of agreement among all reads for a given position, provides a more reliable metric for base-level methylation. We defined all the positions as methylated with a *Percent Modified* value above 0.5. This threshold was determined by calculating the average *Percent Modified* value of methylated bases within motifs identified in our dataset but already known and validated in the literature (see **Supplementary Figure S1**).

All the steps described above are integrated into a custom Nextflow^44^ pipeline, publicly available on GitHub, which automates preprocessing, alignment, methylation analysis, and motif detection^45^. We ran our pipeline in version 0.0.2 and as described in the GitHub manual.

### Integrating methylation detection and analysis with MicrobeMod

We used MicrobeMod on all samples, running both available pipelines: one extracted all RM genes along with their associated motifs from the literature, while the other used STREME^46^ on the Modkit output to identify motifs. MicrobeMod was run with default parameters for both cases.

For isolate LM46, we observed a higher number of 4mC bases not associated with a known motif and a similar pattern for isolate LM41 regarding 5mC. To investigate this observation further, we ran MicrobeMod with a lower *percent_methylation_cutoff*, which adjusts the fraction of methylated reads required to classify a site as methylated. This analysis identified two additional motifs: 5’-TGG-**4mC**-CA-3’ and 5’-T-**5mC**-GA-3’. These motifs were also confirmed using Modkit’s *motif evaluation* function, which allows for assessing specific motifs after the refinement of thresholds for the fraction of modified bases.

## Results

### Detected methylation motifs and sites using MicrobeMod and Modkit

We applied MicrobeMod and Modkit (implemented in our Nextflow pipeline) on the re-basecalled and re-assembled data of the twelve bacterial species (see **Methods**). Next, we summarized the detected motifs by combining the results from both tools (**Table 1)**. In most cases, both tools detected the same motifs (see **Supplementary Table S3** for a comparison of motifs individually identified by each tool). However, we detected a few discrepancies worth mentioning: 1) SA67 motifs: Modkit detected two motifs, CCAYNNNNNNRTC and CCAYNNNNNNTTYG, while MicrobeMod identified a broader, less specific motif, CCAYNNNNNNDTY. This motif spans the two detected by Modkit but with less consistency, suggesting some limitations in MicrobeMod’s ability to distinguish between closely related patterns. 2) MicrobeMod did not detect the motifs GAAANNNNNNGGG in KP13 and GAAGAC in LM46, and Modkit did not detect the motif GAAGNNNNNTAC in SA67. 3) CAGDAC vs. CAGNAC in EF35: MicrobeMod identified CAGNAC, treating the “D” (A or G or T) as an “N” (any base), resulting in a less specific motif with a lower methylation percentage. Modkit identified CAGDAC, which is likely the more accurate representation of the motif.

**Table 1.** List of motifs detected using Modkit and MicrobeMod. The table combines results from both tools and categorizes motifs by the type of methylated base detected: 6mA, 5mC, or 4mC. The methylated base is highlighted in bold. An asterisk is appended if the base is methylated on the complementary strand (e.g., T* and G*). Two motifs, GAAANNNNNNGGG in KP13 and GAAGAC in LM46, present methylation with both 6mA and 4mC. Please note that the sample IDs are the same as those used in the original study^23^.

| Sample ID | Species | Detected motifs with 6mA | Detected motifs with 5mC | Detected motifs with 4mC |
| --- | --- | --- | --- | --- |
| KP04 | *Klebsiella pneumoniae* | GAT*C | CCWG*G |  |
| KP02 | *Klebsiella pneumoniae* | GAT*C,<br>GACNNNNNNGT*C | CCWG*G |  |
| KP13 | *Klebsiella pneumoniae* | GAT*C,<br>AGCNNNNNCT*TC,<br>GAAANNNNNNGGG | CCWG*G | GAAANNNNNNGG*G |
| LM46 | *Listeria monocytogenes* | GAAGAC |  | GAAG*AC,<br>TGG*CCA |
| LM41 | *Listeria monocytogenes* | GAAYNNNNNGT*C | TCG*A |  |
| LM54 | *Listeria monocytogenes* |  |  |  |
| EF35 | *Enterococcus faecium* | CAGDAC |  |  |
| EF26 | *Enterococcus faecium* | CYAANNNNNNGRT*Y |  |  |
| EF22 | *Enterococcus faecium* | CYAANNNNNNGRT*Y |  |  |
| SA63 | *Staphylococcus aureus* | GWAGNNNNNGAT*,<br>CGANNNNNNNT*CC |  |  |
| SA67 | *Staphylococcus aureus* | GAAGNNNNNT*AC,<br>CCAYNNNNNNRT*C,<br>CCAYNNNNNNT*TYG |  |  |
| SA62 | *Staphylococcus aureus* | CCAYNNNNNNT*GT,<br>AGGNNNNNGAT* |  |  |

Our results were consistent across the three replicates from the different labs, and the motifs identified were found to be methylated in over 95–99% of their occurrences across the respective genome. However, two motifs showed lower methylation levels: TGGCCA in LM46 (∼90%) and TCGA in LM41 (∼30%) (**Table 1**). Overall, Modkit provides more comprehensive results (e.g., detecting more motifs or identifying them with higher specificity, such as CAGDAC), although the tool did not detect the motif GAAGNNNNNTAC in SA67. This is likely due to its low occurrence frequency in the genome (only 200 instances).

**Supplementary Table S3** also includes motifs identified using MicrobeMod’s annotation pipeline, which predicts motifs based on methyltransferase genes annotated in the reference genome using the REBASE database. These motifs are thus directly associated with detected methyltransferases in the dataset.

Overall, the annotation pipeline results were broadly consistent with the detected motifs. Still, there were exceptions: 1) EF35: The motif CYAANNNNNNGRTY was absent in the detected data and replaced by the motif CAGDAC. 2) KP13: An additional motif, GAAANNNNNNGGG, was detected *de novo* but not predicted by the annotation pipeline. 3) SA67: The annotation pipeline predicted the motif CCAYNNNNNNTTYG, which was confirmed as methylated. However, another similar motif, CCAYNNNNNNRTC, was detected but not found by the annotation pipeline.

These findings highlight the growing capability of new tools and models to confirm previously known motifs and detect *de novo* motifs that were previously underrepresented or undescribed while also highlighting potential differences.

#### Motif detection accuracy

The methylated bases within the detected 6mA motifs generally exhibited an average *Percent Modified* value ranging from 70 to 90% (see **Figure S1**). Interestingly, the motif GAAANNNNNNGGG from KP13, which comprises both 6mA and 4mC signals, shows a comparably high *Percent Modified* value for 4mC. An outlier was the motif GAAGAC from LM46, where the *Percent Modified* value for both 6mA and 4mC was lower than for all other samples and motifs and only ranged from 55 to 60%. The 5mC motif CCWGG, detected across all three *K. pneumoniae* isolates, exhibited even higher *Percent Modified* values, ranging from 85 to 95% in all isolates and replicates, except for KP02 from LAB3, which showed a notably lower value of 65%. This lower value could be attributed to the reduced coverage of this isolate from LAB3, which may have led to a lower *Percent Modified* value and an increased number of unmethylated occurrences, which were not confirmed by other replicates with higher coverage (see **STab 1** for coverage information).

Additionally, the motifs TCGA (5mC) and TGGCCA (4mC), which reported lower percentages of methylated occurrences across the genome (< 95%), also exhibited much lower *Percent Modified* value, around 30%.

Overall, using our approach, up to 95% of 6mA bases (with *Percent Modified* > 0.5) were found within a motif (see **Figure S1**). The only exception is one isolate from *Listeria* where no motifs were detected (LM54). In this isolate, almost no mutations with a high *Percent Modified* base were detected (> 0.7)

By applying less stringent parameters in MicrobeMod (see **Methods** for details) to analyze samples LM41 and LM46, which appeared to have higher levels of 5mC and 4mC, respectively, we identified two additional motifs: TCGA (5mC) and TGGCCA (4mC). As a result, the 5mC and 4mC modifications were explained in up to 90% of the bases within the motifs. However, we detected more 4mC bases in *Klebsiella* isolates, but these were not associated with a specific motif. We further observed that these modified bases clustered near GATC or CCWGG motifs, raising the possibility that these are either genuine modified bases or technical artifacts that still need further clarification.

### Analysis of hypermethylated genes and motifs in regulatory regions

We found certain regions of the genomes to be enriched in methylation. Using the genome annotation obtained with Bakta, we calculated the methylation density for each gene by dividing the number of methylated bases (*Percent Modified* > 0.5) by the gene size. Only the LAB3 dataset results are shown for this analysis, as the results were consistent between the three replicates. The results are presented as violin plots, illustrating the distribution of methylation density for each gene, grouped by COG (clusters of orthologous groups) category based on the Bakta annotation (see **Figure 1**, **Figures S2** and **S3**).

**Figure 1.**
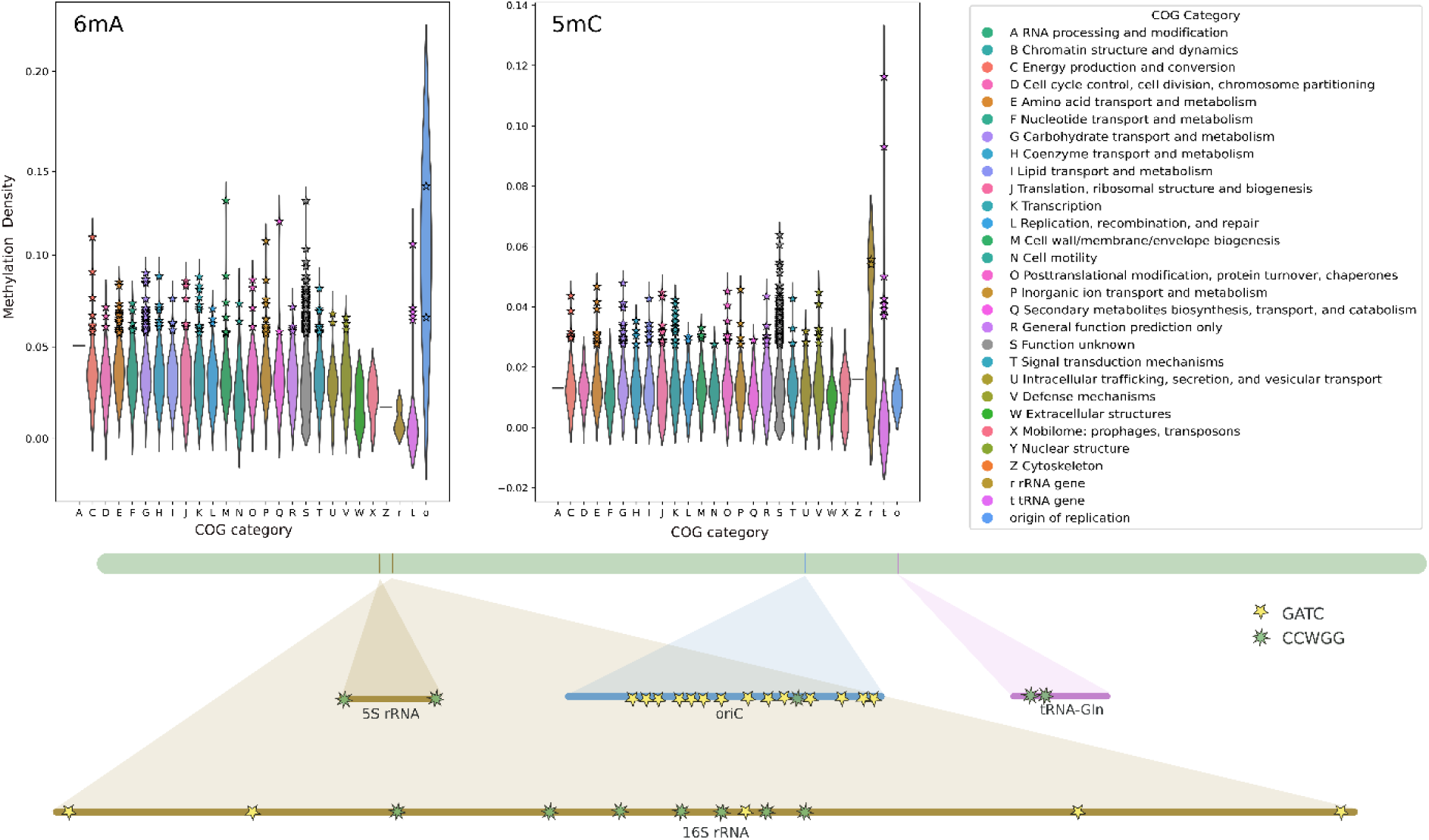
Methylation density of the genes of the KP04 isolate for 6mA (top left) and 5mC (top center). The top 5% of genes with the highest methylation density are highlighted with a star. On the bottom left, the hypermethylated genes are shown on the genome of KP04. COG - clusters of orthologous groups derived from the Bakta annotation.

For *Klebsiella pneumoniae,* we conducted this analysis for both 6mA and 5mC modifications since both bases were extensively present in the data. This is consistent with the methylated motifs GATC and CCWGG, respectively, which are short, palindromic, and persistently distributed across the genome. **Figure 1** exemplarily shows the methylation density of the genes of the isolate KP04 for 6mA and 5mC. In addition to COG categories, we included methylation levels for the origin of replication (*ori*), given the well-established role of the GATC motif in initiating the cell cycle and its high abundance in this region^47,48^. We also included rRNA and tRNA genes in the analysis, as an initial rapid inspection of the data suggested increased methylation levels in these regions.

Our analysis confirmed that the *ori* region of KP04 is enriched in 6mA bases (**Figure 1**), with no other category standing out, except for a few genes displaying similar methylation levels to the *ori*. In particular, one such gene belongs to the COG category *Cell wall/membrane/envelope biogenesis*. It is also worth mentioning that many genes fall under the COG classification S = *Function unknown*.

The plot for 5mC (**Figure 1**, center) revealed that rRNA and tRNA genes are enriched in 5mC bases, whereas most other categories did not exhibit significant enrichment. To further investigate, we visualized these patterns using IGV^49^, confirming that these regions were indeed enriched with either GATC or CCWGG motifs (bottom part of **Figure 1**). In some cases, such as for 5S rRNA and 16S rRNA, the motifs flanked the genes at both the 5’ start and 3’ end. However, this pattern was inconsistent across all rRNA genes; for instance, 23S rRNA displayed a CCWGG motif only at the 5’ start of the gene. Despite their small size, tRNA genes often contained one occurrence of the CCWGG motif, with two occurrences found specifically in glutamine tRNA genes. The same patterns were observed in the other two isolates of *Klebsiella pneumoniae* (see **Figure S2)**. For the other species, only 6mA methylation was analyzed (see **Figure S3**), but similar trends emerged. These included rRNA and tRNA genes exhibiting high methylation density and genes in the COG category J = *Translation, Ribosomal Structure, and Biogenesis*.

We further explored methylation patterns by directly examining the *Percent Modified* values for each base rather than applying a threshold to define bases as methylated. We computed a methylation density by averaging these raw *Percent Modified* values in windows of 1000 bases, and we aligned these values with genome annotations. In *Staphylococcus aureus,* this analysis revealed higher methylation values associated with rRNA genes (**Figure 2**).

**Figure 2.**
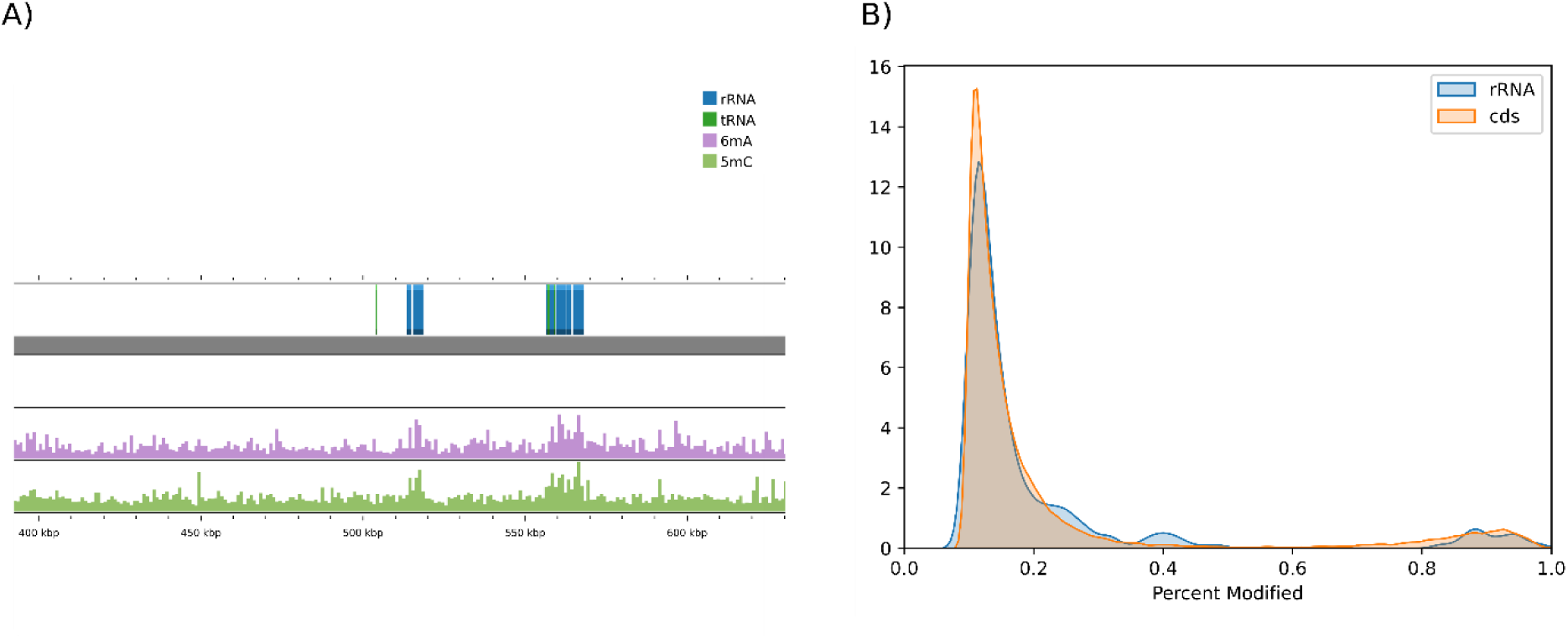
Average methylation density in SA63 alongside rRNA and tRNA genes (A) and distribution of *Percent Modified* in SA63 for coding DNA sequences (CDS) and rRNA genes (B). On the left, methylation density was calculated using 1000-nucleotide windows and aligned with the annotation from Bakta. We observed a higher methylation density corresponding to the rRNA and tRNA genes for both 6mA and 5mC. On the right, we found that rRNA genes have a higher number of bases with a *Percent Modified* between 0.2 and 0.4, which may contribute to the observed increase in methylation density.

When plotting the distribution of *Percent Modified* values (ranging from 0.1 to 1), rRNA genes in *S. aureus* showed a higher proportion of bases falling in the range of 0.2 to 0.4. Unlike *K. pneumoniae*, where the CCWGG motif was enriched in these regions, no specific motif was associated with methylation in *S. aureus*. Instead, the data highlighted a broader pattern of mid-to-low methylation levels (< 0.5) across rRNA genes.

We investigated whether specific motifs were enriched in the promoter regions of genes or at their start/end positions. For this analysis, we defined the promoter region as the 40 bases upstream of the gene, the start of the gene as the first 40 bases following the transcription start site, and the end of the gene as the last 40 bases. The choice of 40 bases was made to capture the bacterial core promoter, including the −10 and −35 elements^50^, and was applied equally to the start and end regions. For each motif, we calculated the percentage of occurrences within these three regions (**Figure 3**). Based on the annotation from Bakta, we estimated that 10.5% - 11% of the genome falls within one of these categories, and our results suggest that only a few motifs are more concentrated in these regions than expected. Examples include the motif AGGNNNNNGAT from SA62, the motif CAGDAC from EF35, and the motif GAAANNNNNNGGG from KP13, which were more frequently found in these regions than expected. The motif CCAYNNNNNNRTC in SA67 is also more frequent in the promoter regions. In contrast, the motif TGGCCA was least frequently observed in promoter regions.

**Figure 3.**
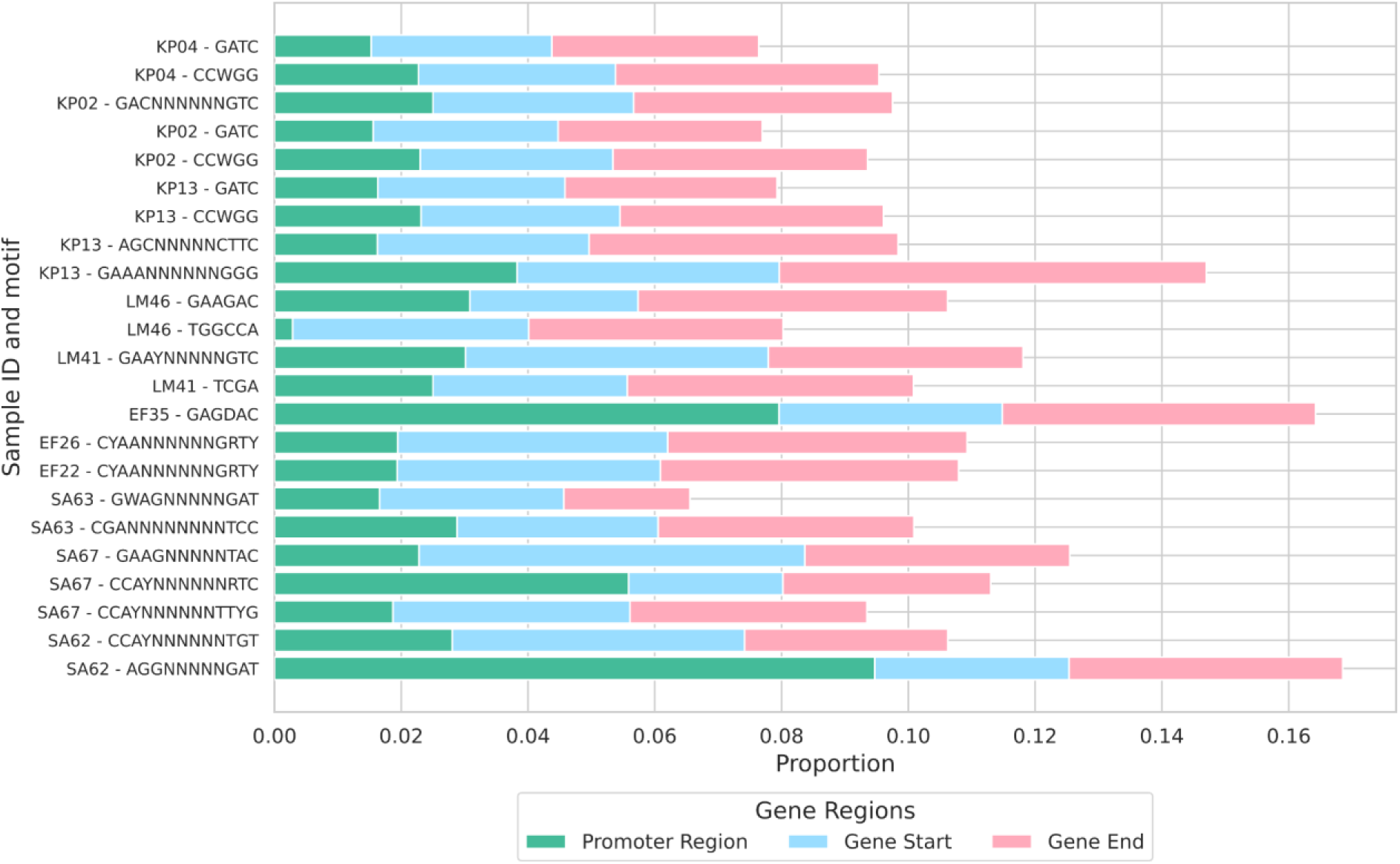
The proportion of methylated bases in promoter regions, gene start, and gene end for each detected motif. The promoter region is defined as the 40 bases upstream of the gene, the gene start as the first 40 bases following the transcription start site, and the gene end as the last 40 bases of the gene.

### Assessment of potentially methylation-induced basecalling errors

In nanopore sequencing, the raw signal is generated by overlapping k-mers of nucleotides passing through the nanopore rather than by individual bases. This process can sometimes lead to inaccurate basecalling, particularly in regions surrounding modifications^21,24^. However, the introduction of newer basecalling models has potentially mitigated this issue. To evaluate the current extent of these improvements, we compared reads basecalled with 2023-09-22_bacterial-methylation^51^ to those basecalled with the latest tool+model combination: Dorado v0.8.1 and basecalling model SUP v5.0.0.

#### Analysis of R and Y methylation-induced ambiguous positions

Previous studies have found that it was a particular challenge for the basecaller to distinguish between G and A (producing ambiguous bases denoted as R in the IUPAC system) or between C and T (Y in the IUPAC system)^23,52^. Our results confirm these findings. By comparing ambiguous positions with methylated data obtained from Modkit, we found that these ambiguities are linked to a methylated adenine located at the position following the error in the 5’ direction. More specifically, an ambiguous position R (G or A) is found immediately before a methylated adenine, while an ambiguous position Y (C or T) corresponds to a G on the reverse strand preceding a methylated adenine. Thus, these two scenarios are equivalent, differing only in strand orientation.

We then used the pipeline MPOA^24^, which identifies and masks ambiguous positions independently of any methylation calling, to evaluate whether the frequency of R and Y bases was reduced in reads basecalled using the new Dorado model. We observed a noticeable reduction in ambiguous R and Y bases using the SUP v5.0.0 basecalling model in samples previously classified as ‘problematic’ based on a comparison to short-read data^23^ **(Figure S4)**. As expected, two samples, LM54 and EF35, classified as not problematic in the original study (see list of the original problematic isolates in **Table S1**), showed no or virtually no ambiguous bases with the old and new models. Meanwhile, two previously non-problematic isolates, KP13 and SA62, now had ambiguous bases, but without a clear bias toward R and Y bases, as usually observed in problematic strains. Interestingly, KP04, classified initially as problematic and still after re-sequencing with the latest translocation speed of 400 bp/s (see **Supplementary Table S1**), did not exhibit a bias for R and Y bases, suggesting that these ambiguous bases were not methylation-induced. Lastly, KP02 and LM46 showed no reduction in the number of R and Y bases, indicating that the basecaller still struggles with these samples despite improvements in the new models.

#### Repurposing Hammerhead to evaluate strand-specific basecalling errors

Many ambiguous positions observed result from strand-specific error patterns near the modification sites. Specifically, the strand carrying the modification tends to be less accurate, and basecalling can result in a different base than the non-modified strand^24^ (in our case, an A instead of a G preceding 6mA). The tool Hammerhead^52^ leverages this error pattern to detect modifications directly from the FASTQ. We repurposed Hammerhead to assess whether improvements in the basecalling models would reduce the number of “potential modification sites” identified by this method.

As expected, due to the improved accuracy of the v5 basecalling models, our results show a remarkable decrease in modification sites identified by the Hammerhead approach for most samples (see **Figure S5**). In many cases, Hammerhead detected only a few sites, except for KP02, LM46, and SA67, which still presented identifiable sites.

#### DNA logo visualization

MPOA allows visualization of the regions surrounding ambiguous positions as a DNA logo. We collected all logos for R and Y bases, showing how these ambiguous positions align with our identified DNA motifs (see **Figure S6**). In most cases, one detected motif is present in the DNA logo. **Figure 4** compares previous^23^ and new (v5 model) basecalling results from MPOA, Hammerhead, and the DNA logos for the isolates most affected by strain-specific methylation-induced errors.

**Figure 4.**
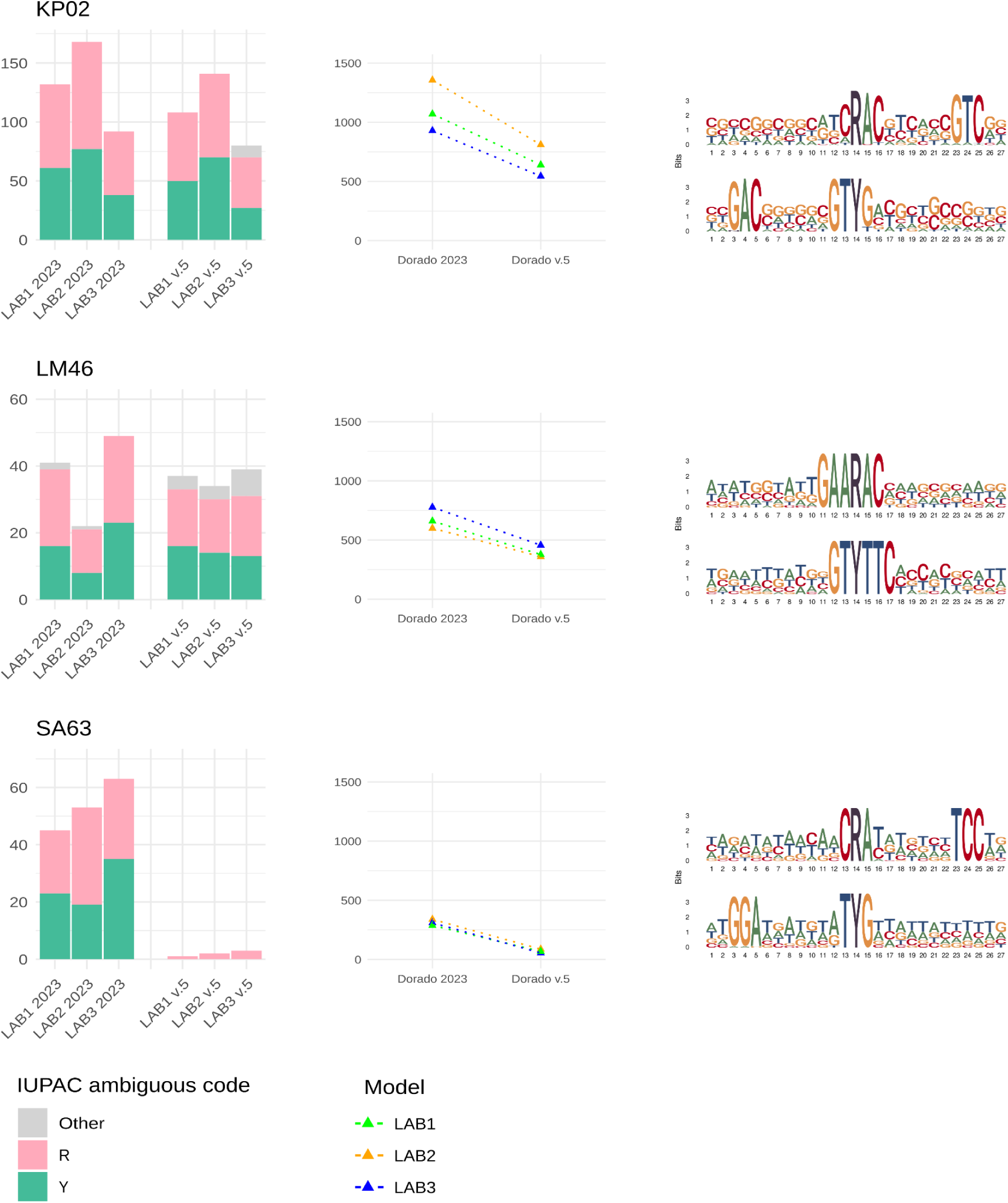
Comparison of Dorado models 2023-09-22_bacterial-methylation (2023) and v5.0.0 using MPOA and Hammerhead for isolates KP02, LM46, and SA63. The number of ambiguous positions (R and Y) is substantially lower in SA63, while KP02 and LM46 show similar values (left column). We use Hammerhead to identify and compare the number of potentially methylated sites between the new and the old model (center column). On the right, DNA motif logos for the positions surrounding R and Y are obtained from MPOA using the older model based on the LAB3 dataset, clearly showing that the ambiguous positions R and Y correspond to methylation motifs found to be active in these isolates.

## Discussion

### Nanopore sequencing and associated methods allow the comprehensive detection of bacteria methylation motifs and sites

In this study, we comprehensively analyzed the methylation profiles of four public health-relevant bacterial species using nanopore sequencing and currently available software tools. We re-analyzed raw signal data from a previous study and presented robust results using triplicates, the latest nanopore chemistry, and R10.4.1 flow cells. Our results revealed distinct species-specific DNA modification patterns reproducible between replicates, including methylated motifs and heavily methylated genomic regions.

A key question addressed in our work was whether current computational tools enable reliable detection of known and *de novo* methylation motifs. Using the tool MicrobeMod and our custom Nextflow pipeline, which incorporates ONT’s Modkit tool, we identified known motifs validated through cross-referencing with the REBASE restriction-modification database. Additionally, we detected several novel motifs with high confidence, characterized by methylation in more than 95% of motif occurrences and consistent methylation across reads, as indicated by high *Percent Modified* scores (ranging from 70% to 90%). However, two motifs showed a higher proportion of unmethylated occurrences and lower *Percent Modified* scores, around 30%. These cases require further investigation to determine whether the bases are truly modified and, if so, to understand why some instances remain unmethylated.

By applying a threshold of 50% for the *Percent Modified* score to classify a base as methylated, we could explain up to 95% of the modified bases through motifs. We observed an exception to this trend in the 4mC bases of *Klebsiella pneumoniae*, which were not directly located within specific motifs, but were consistently clustered near GATC and CCWGG sequences in all KP strains and their replicates. This observation raises the question of whether these C-bases are really 4mC-modified or just wrongly classified as 4mC, since the signal actually originates from 6mA (GATC motif) or 5mC (CCWGG) bases.

Based on our results, we conclude that state-of-the-art nanopore sequencing enables the reliable identification of motifs that are consistently methylated throughout the bacteria genome. This includes short motifs, such as palindromic sequences from Type II RM systems, short non-palindromic motifs associated with Type III RM systems, and longer, more complex motifs characteristic of Type I RM systems. However, challenges persist in identifying motifs that exhibit inconsistent methylation or occur at lower frequencies. In one case (GWAGNNNNNGAT in SA67), for instance, Modkit failed to recognize a motif with low frequency (around 200 occurrences throughout the genome). Overall, the results from Modkit appeared more complete than those from MicrobeMod; combining data from both methods yielded the most reliable results.

An interesting feature observed in our motifs table is that all motifs, except CAGDAC in EF35, were methylated on both strands. This was expected for palindromic sequences since the sequence is identical on both strands, resulting in the same position being methylated both on the reverse and forward strands. However, we also observed this for longer Type I bipartite motifs, where different parts of the motif were methylated on opposite strands. In the case of GAAANNNNNNGGG in KP13, the adenine at position 4 is methylated, while the guanine at position 12 corresponds to a methylated cytosine (4mC) on the reverse strand. We also observed this combination of distinct methylation types (6mA and 4mC) in GAAGAC in LM46, where the methylated bases were adjacent: position 4 (4mC on the reverse strand) and position 5 (6mA on the forward strand).

Given these results, a potential idea for a future tool could involve implementing an algorithm that scans for motifs by simultaneously detecting methylation on both strands rather than analyzing each modification individually. Such an approach could enhance the accuracy of motif identification, especially in cases where one modification exhibits a lower percentage of methylation.

Our analysis of hypermethylated regions and genes showed that rRNA genes are enriched in CCWGG motifs, consistent with previous findings^13,53^. Additionally, we detected one or two occurrences of CCWGG motifs in many tRNA genes, which has also been noticed previously^54^. Furthermore, we noted an overall higher density of CCWGG in genes associated with the COG category “*Translation, Ribosomal Structure, and Biogenesis*”. Interestingly, in rRNA genes from other species, particularly *Staphylococcus aureus*, we noted an increase in positions with a low *Percent Modified* score for 5mC (>0.2 and <0.5), indicative of incomplete methylation of reads, despite the absence of specific detectable motifs. These bases with low modification levels require further investigation, as the low *Percent Modified* score could be due to technical issues.

We conclude that bacterial DNA methylation influences the regulation of ribosomal function and protein synthesis, potentially impacting cellular processes such as growth and proliferation. Thus, we provide further evidence that methylation is essential not only in restriction-modification systems but also plays a broader role in the epigenetics of the cell. Our analysis supported these findings by identifying a subset of motifs that are more present than expected in promoter regions and/or at the beginning or end of genes.

### Recent basecalling models reduce strand-specific ambiguity, but errors still persist in certain motifs, affecting reproducibility

Our analysis is based on a dataset including three replicates from different institutes. This setup allowed us to compare results from three independent sequencing runs and evaluate the reproducibility of methylation analysis using the latest nanopore technologies.

The results were largely consistent across replicates, with the same motifs detected in all cases. However, a few isolates showed more unmethylated occurrences for specific motifs. These discrepancies could be attributed to lower sequencing coverage and reduced accuracy in these isolates, as in the case of CCWGG in KP02 from LAB3, where about 1.6% of the occurrences were unmethylated. Similarly, we observed slight increases in unmethylated occurrences in other low-coverage isolates (e.g., EF35 and EF22 from LAB3), while the motifs were almost fully methylated in the other two labs. One hypothesis, to be further tested, entails that specific motifs may exhibit varying methylation levels, which could fluctuate during cultivation or under different conditions and should be investigated further.

Many bioinformatics tools are available for extracting methylation levels from bacterial genomes^28,55–57^. For our analysis, we chose to compare and integrate results from MicrobeMod, specialized for bacteria, and a custom pipeline that includes ONT’s Modkit tool, as both are compatible with the latest flowcell version (R10.4.1) and the recently released POD5 format for raw Nanopore signal data. Many previous tools were developed for the outdated R9 flowcells and not updated accordingly or are more specifically focused on eukaryotic (mainly human) DNA methylations.

We have made our pipeline publicly available, enabling users to run Modkit efficiently on raw signal POD5 data and extract subsets of bases identified as methylated. We plan to extend this work in the future by incorporating new features, such as a more comprehensive overview of the results and improved comparison with genome annotations. By that, we also aim to scan for antibiotic resistance genes and measure their methylation levels.

Recent advancements in methylation detection using nanopore sequencing have significantly improved the accuracy of detecting methylated bases and the basecalling of nearby positions. The latest v5 models, with SUP and HAC modes, allow for directly detecting 6mA, 5mC, and 4mC bases. Despite this progress, challenges with basecalling near methylated sites remain. Specifically, we observed that the basecaller often misclassified a guanine preceding a methylated adenine as an adenine. This led to ambiguous sites where reads were inconsistent, with some indicating a guanine and others an adenine, resulting in R (mixed bases) according to the IUPAC code. These errors are particularly evident for a subset of motifs (see **Figures S4** and **S6**). For example, in Biggel *et al*.^22^, the GAAGAC motif was associated with erroneous sites in a subset of *Listeria monocytogenes* isolates with this actively methylated motif. We detected this motif also in isolate LM46, and, along with the motif GACNNNNNNGTC in KP02, it remains problematic in our dataset regarding wrong basecalls. The reasons why specific motifs are more difficult for the basecaller are still unclear. One possible explanation lies in the underlying data used to train the models, which might be geared towards certain types of bacteria.

In our dataset, the issue seems to be sequence context-specific: sequence logos of these problematic positions (**Figure S6**) reveal that a cytosine often precedes the ambiguous sites, and the methylated adenine is frequently followed by either a cytosine or a thymine, resulting in the error-prone motif CRAY (where Y is either C or T). In the case of GAAGAC, it may also play a role that the ambiguously basecalled G at position 4 is methylated on the opposite strand (4mC).

These basecalling errors can be directly linked to methylation, as the issues occur only on the methylated strand. The non-methylated strand, in contrast, remains accurate in comparison to short-read data. One potential solution is to focus on positions with high differences between the two strands and use information from the non-methylated strand for better accuracy. However, this requires previous knowledge regarding which strand is methylated or which motifs are actively present in the genome. Another way to investigate the interaction between methylation and base calling errors is to perform additional PCR amplification before nanopore sequencing to obtain a purer signal without interfering modifications.

Hammerhead is a tool that uses these strand differences to infer potential methylation sites and is particularly valuable for users who only have FASTQ files rather than raw signal data. However, our results show that such errors occur less frequently with the newer v5 models, such errors are becoming less common. Using Hammerhead with standard pipeline settings, we found very few cases of potential modification in multiple isolates, suggesting that this approach may not be as successful when basecalling accuracy for modified bases improves.

However, Hammerhead could still be useful for detecting unknown, rare, or less-characterized motifs contributing to basecall errors, especially in underrepresented bacterial species with limited or no training data integrated into the basecalling model. These errors may serve as valuable features for identifying potential modifications in such cases.

## Conclusions

Our study represents a genome-wide investigation of the methylation profiles of four bacterial species relevant to public health, using three isolates each. We compared raw signal data from three independent sequencing runs by combining the DNA sequencing results from three laboratories. To address the need for a comprehensive benchmark of the latest methylation tools available for microbial nanopore data, we evaluated the reproducibility of methylome profiling in selected bacteria. While our study provides valuable insights, it is still necessary to continuously benchmark the latest tools and sequencing advancements. However, the lack of available nanopore raw signal data also challenges comprehensive benchmarks and further developments. Due to their size, the frequently changing specifications of the file format, and the lack of supporting online repositories and guidelines, they cannot be exchanged as straightforwardly as FASTQ files. Nevertheless, the recent advancements in nanopore technology, which is especially well-suited for bacterial genome sequencing and reconstruction, are opening new opportunities for in-depth analysis of the bacterial methylome, ultimately helping to decipher the mechanisms underlying the epigenetics of bacteria.

## Data availability

The sequencing data used in this analysis is available from the original study^23^. Our Nextflow pipeline for methylation detection from nanopore data is freely available at https://github.com/rki-mf1/ont-methylation under GPL v3 license. Additional result files, final outputs, and scripts can be found at the Open Science Framework: https://osf.io/jfy75.

## Acknowledgments

We would like to thank the IT team at the RKI and, in particular, the associated access to a high-performance cluster with strong graphics card support. VG thanks the RKI Special Research Funds Program (SOFO) for support.

## Author information

### Contributions

VG and MH conceptualized and designed the study; JDH, CK, and GEW shared data for analysis; VG performed the analyses, analyzed the sequencing data, and created all figures; JDH, ML, CB, CK, and GEW provided valuable feedback during several discussions helping guide the study; MH supervised the study; VG and MH drafted the manuscript. All authors read, commented, and approved the final manuscript.

### Corresponding authors

Valentina Galeone, Martin Hölzer.

## Ethics declarations

### Ethics approval and consent to participate

Not applicable.

### Consent for publication

All authors have read and consented to publish this manuscript.

## Supplementary Material

**Figure S1.**
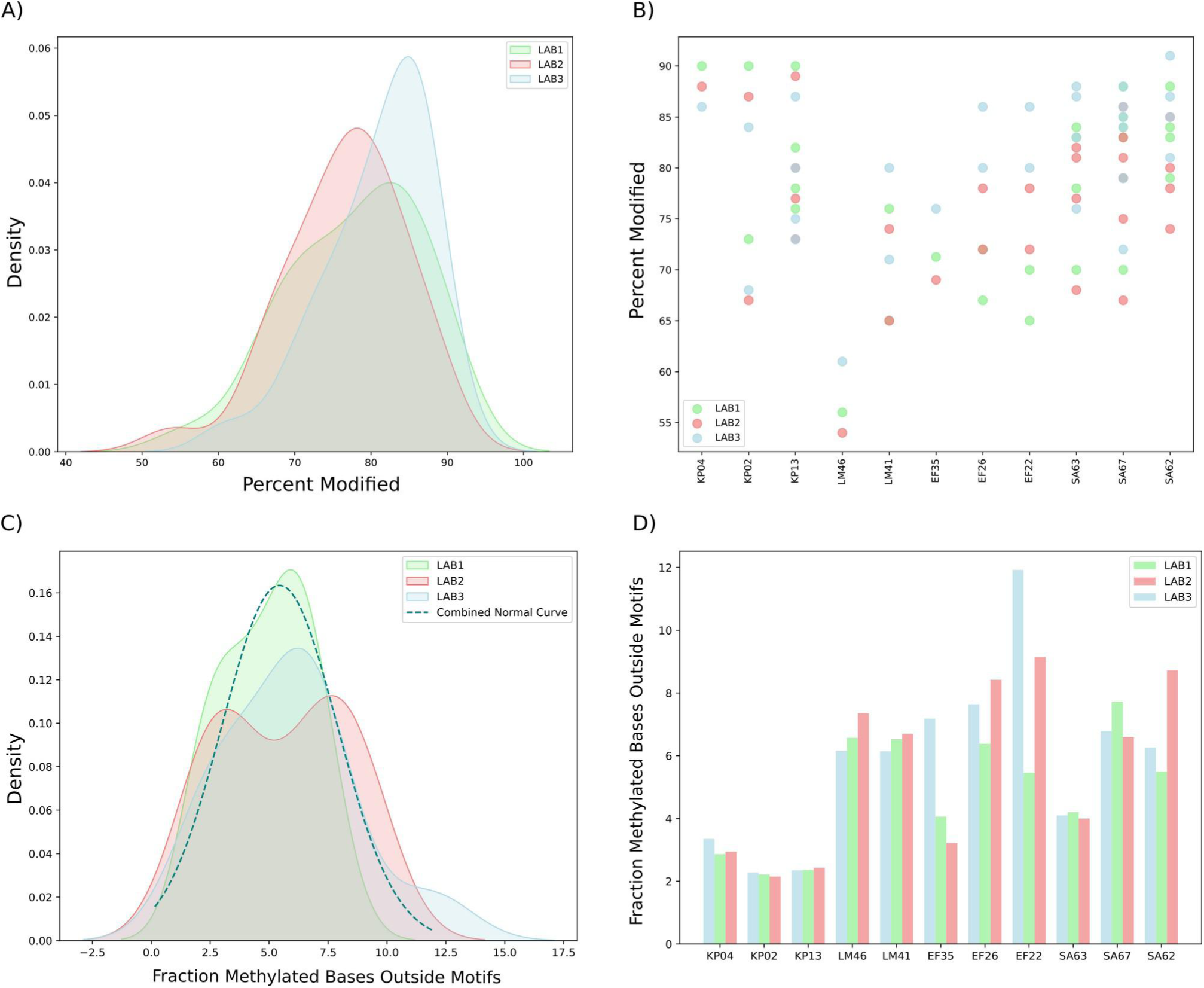
Top panels: Average *Percent Modified* values of methylated bases within detected motifs for 6mA. **(A)** Distribution of *Percent Modified* values for each replicate. **(B)** Distribution of *Percent Modified* values, divided by sample ID. Most motifs were validated from the literature, including the motif in LM46, which shows a lower *Percent Modified* (55-60%). Based on these results, we defined bases with *Percent Modified* > 0.5 as methylated in our study. Bottom panels: Fraction of methylated bases (*Percent Modified* > 0.5) that are not explained by a motif for 6mA. **(C)** Distribution for each replicate. **(D)** Distribution divided by sample ID. These results indicate that approximately 95% of methylated bases are explained by a motif.

**Figure S2.**
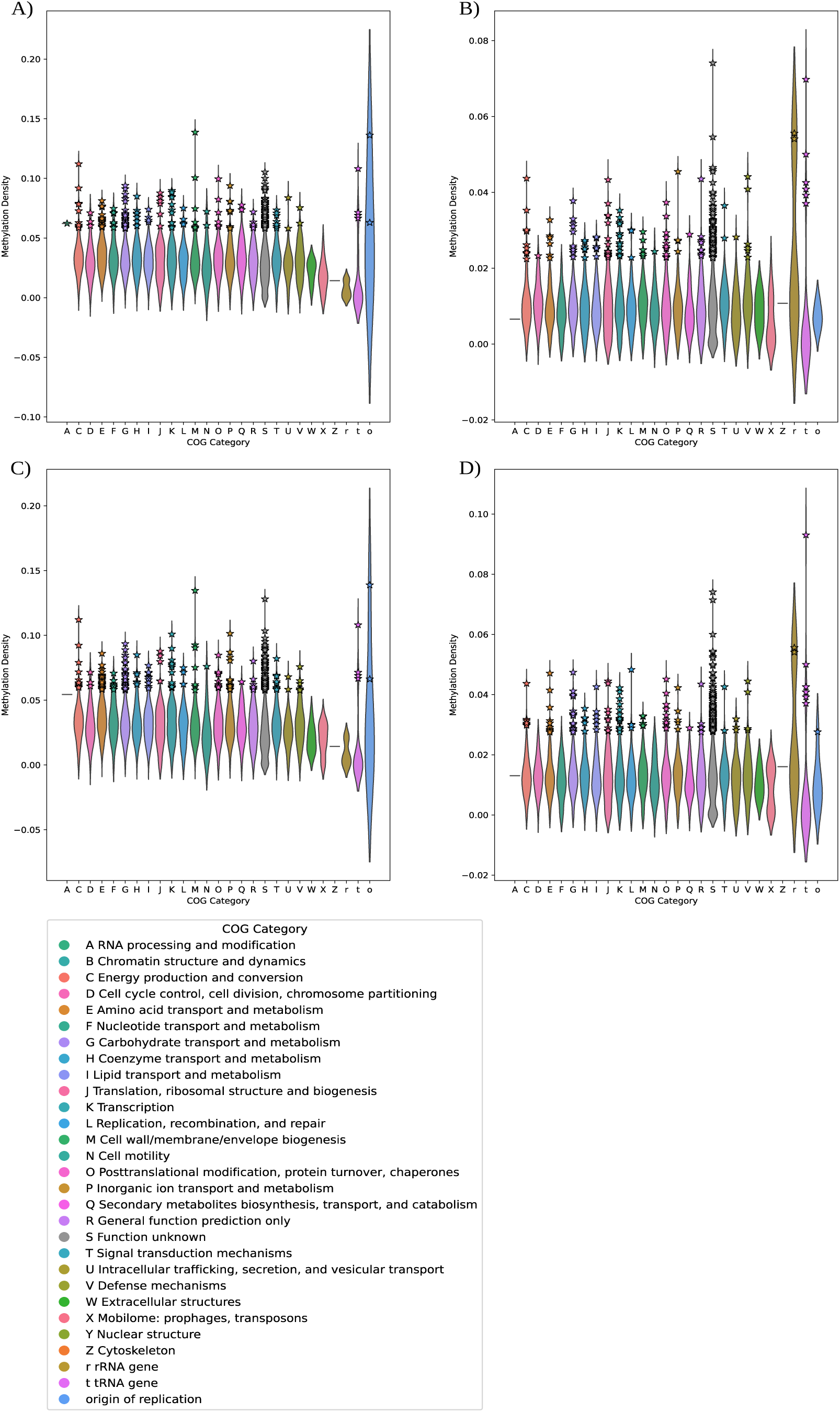
Violin plots of methylation density of the genes of KP02 and KP13 isolates: panels **A)** and **B)** show 6mA and 5mC methylation density for KP02, while panels **C)** and **D)** show 6mA and 5mC for KP13. The genes are grouped by gene function according to the COG classification from Bakta.

**Figure S3.**
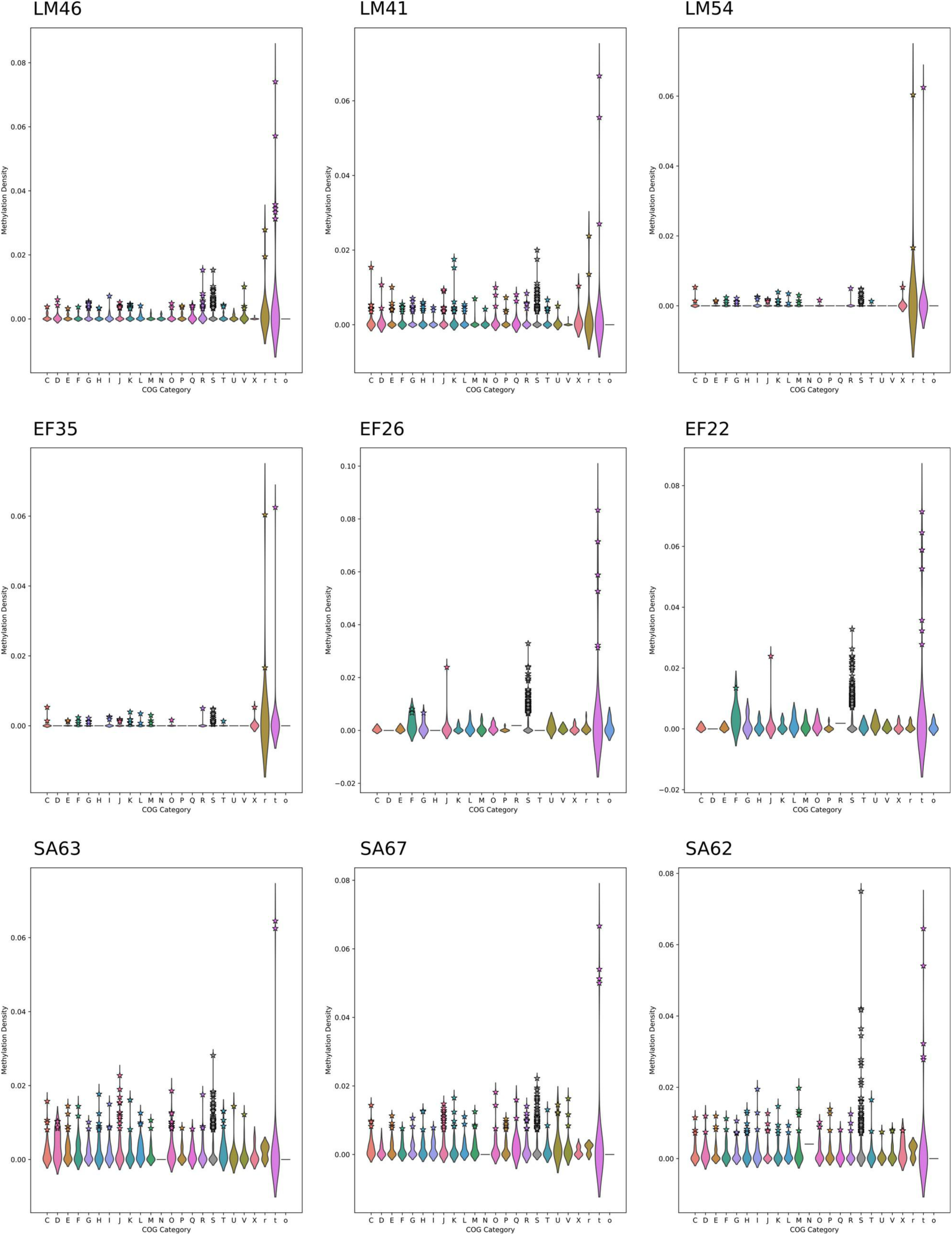
Violin plots of methylation density of the genes of LM, EF, and SA for 6mA. The genes are grouped by gene function according to the COG classification from Bakta (for the color code, please refer to the COG legend in **Supplementary Figure S2**).

**Figure S4.**
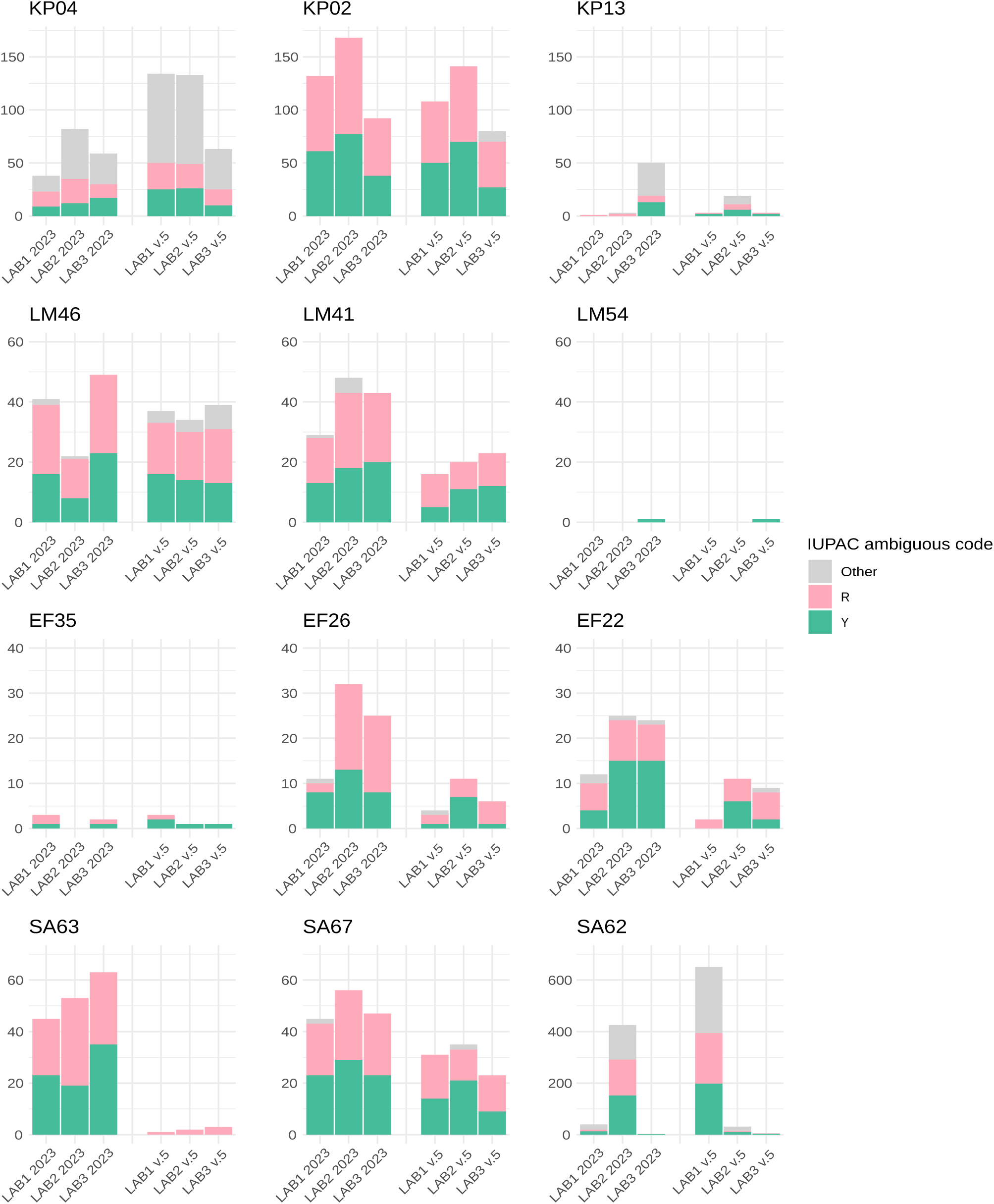
Comparison of ambiguous positions (R = A or G, Y = C or T in IUPAC code) between the old model 2023-09-22_bacterial-methylation (2023) and the new Dorado model v5 across three replicates obtained with the pipeline MPOA. The data show a decrease in the number of ambiguous positions for most samples, with the exception of KP02 and LM46.

**Figure S5.**
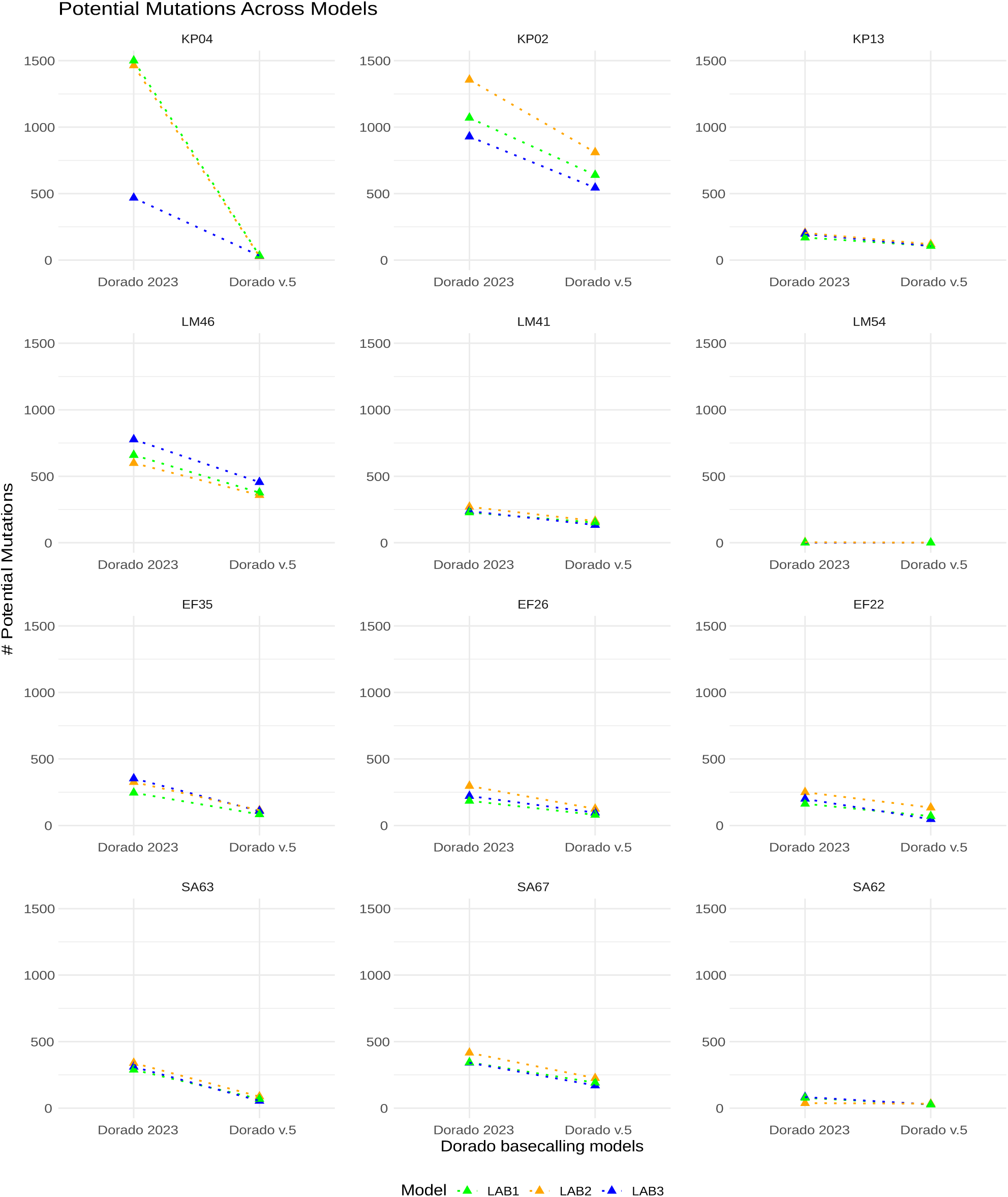
Comparison of the number of potential mutation sites identified by Hammerhead for the old model 2023-09-22_bacterial-methylation (2023) and the new Dorado model v5.

**Figure S6.**
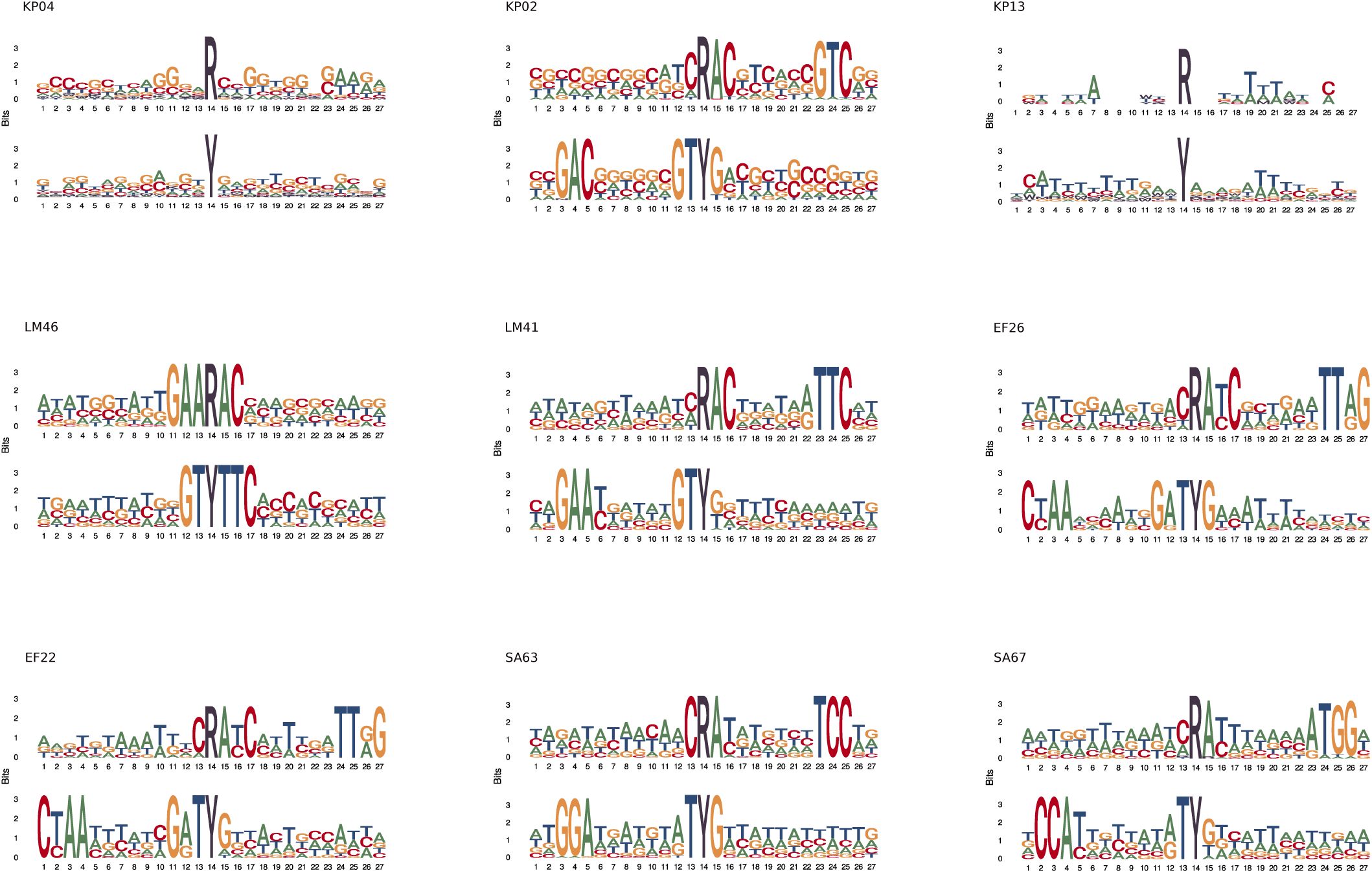
DNA logos generated with MPOA for the strains with R and Y ambiguous positions, using the Dorado basecalling models 2023-09-22_bacterial-methylation on the LAB3 dataset, highlighting the bases surrounding ambiguous positions. In most cases, motifs detected with our pipelines/MicrobeMod are clearly identifiable within the sequences.

**Table S1.** Matching of nanopore sequencing reads (initially sequenced with a translocation speed of 260 bp/s (4 kHz) and later re-sequenced with a translocation speed of 400 bp/s (5 kHz) to Illumina short-read (SR) control data, as reported in the original study (Dabernig-Heinz et al. 2024).

| SAMPLE ID | Matching SR (260 bp/s) | Matching SR (400 bp/s) |
| --- | --- | --- |
| KP04 | No | No |
| KP02 | No | Yes |
| KP13 | Yes | Yes |
| LM46 | No | No |
| LM41 | No | Yes |
| LM54 | Yes | Yes |
| EF35 | Yes | Yes |
| EF26 | No | Yes |
| EF22 | No | Yes |
| SA63 | No | Yes |
| SA67 | No | Yes |
| SA62 | Yes | Yes |

**Table S2.**
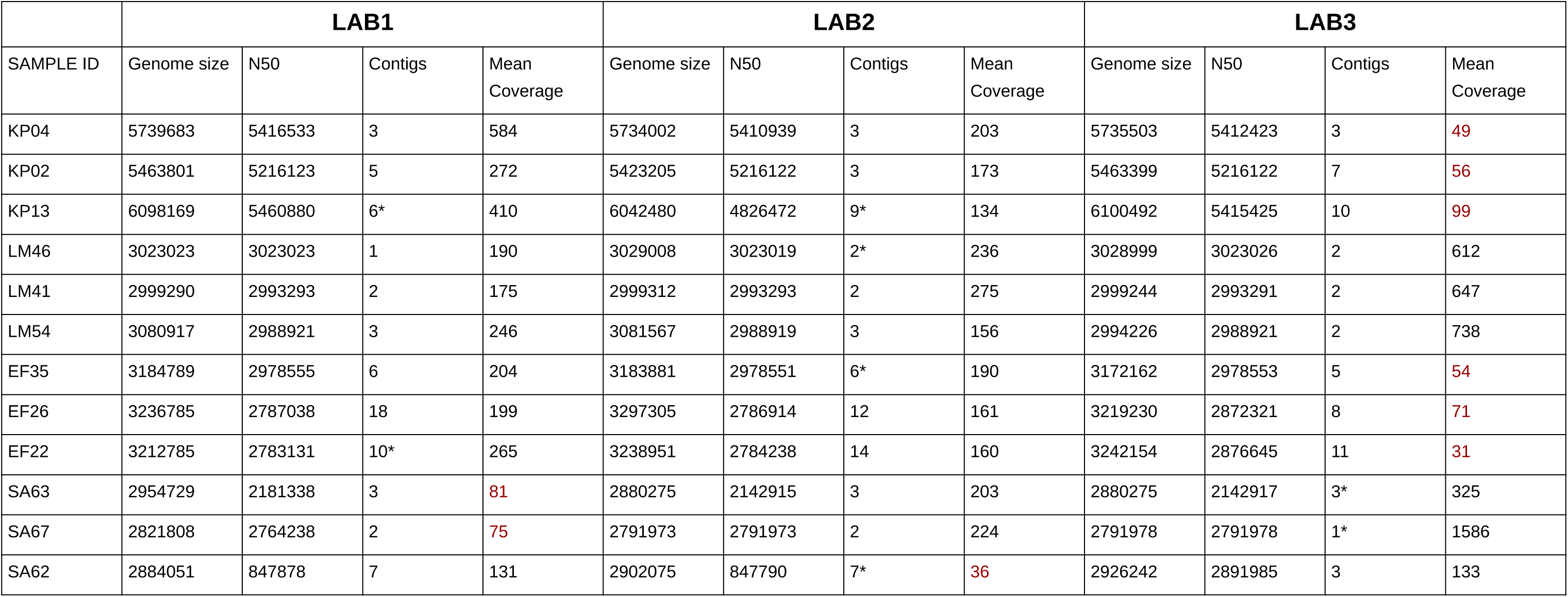
Assembly quality metrics and read coverage statistics for analyzed bacterial genomes. The table includes the total size of each genome assembly, N50 values, the number of contigs, and the mean read coverage. Mean coverage values displayed in red indicate isolates with coverage <100X. A contig number with an asterisk indicates the assemblies that were obtained with the flag “--meta” in flye.

|  | LAB1 |  |  |  | LAB2 |  |  |  | LAB3 |  |  |  |
| --- | --- | --- | --- | --- | --- | --- | --- | --- | --- | --- | --- | --- |
| SAMPLE ID | Genome size | N50 | Contigs | Mean Coverage | Genome size | N50 | Contigs | Mean Coverage | Genome size | N50 | Contigs | Mean Coverage |
| KP04 | 5739683 | 5416533 | 3 | 584 | 5734002 | 5410939 | 3 | 203 | 5735503 | 5412423 | 3 | 49 |
| KP02 | 5463801 | 5216123 | 5 | 272 | 5423205 | 5216122 | 3 | 173 | 5463399 | 5216122 | 7 | 56 |
| KP13 | 6098169 | 5460880 | 6* | 410 | 6042480 | 4826472 | 9* | 134 | 6100492 | 5415425 | 10 | 99 |
| LM46 | 3023023 | 3023023 | 1 | 190 | 3029008 | 3023019 | 2* | 236 | 3028999 | 3023026 | 2 | 612 |
| LM41 | 2999290 | 2993293 | 2 | 175 | 2999312 | 2993293 | 2 | 275 | 2999244 | 2993291 | 2 | 647 |
| LM54 | 3080917 | 2988921 | 3 | 246 | 3081567 | 2988919 | 3 | 156 | 2994226 | 2988921 | 2 | 738 |
| EF35 | 3184789 | 2978555 | 6 | 204 | 3183881 | 2978551 | 6* | 190 | 3172162 | 2978553 | 5 | 54 |
| EF26 | 3236785 | 2787038 | 18 | 199 | 3297305 | 2786914 | 12 | 161 | 3219230 | 2872321 | 8 | 71 |
| EF22 | 3212785 | 2783131 | 10* | 265 | 3238951 | 2784238 | 14 | 160 | 3242154 | 2876645 | 11 | 31 |
| SA63 | 2954729 | 2181338 | 3 | 81 | 2880275 | 2142915 | 3 | 203 | 2880275 | 2142917 | 3* | 325 |
| SA67 | 2821808 | 2764238 | 2 | 75 | 2791973 | 2791973 | 2 | 224 | 2791978 | 2791978 | 1* | 1586 |
| SA62 | 2884051 | 847878 | 7 | 131 | 2902075 | 847790 | 7* | 36 | 2926242 | 2891985 | 3 | 133 |

**Table S3.**
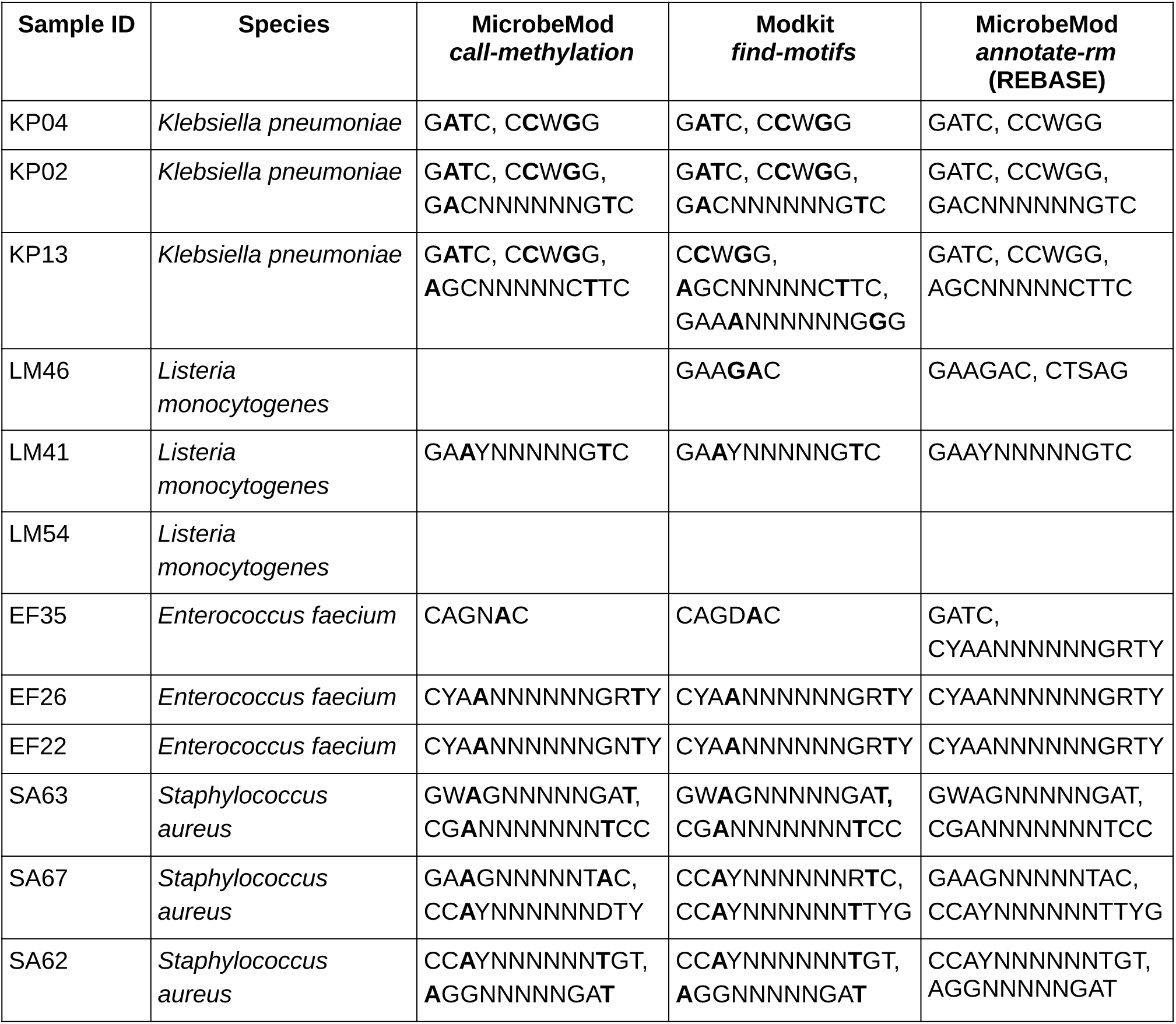
Comparison of methylated motifs identified by MicrobeMod’s call-methylation pipeline, Modkit (find-motifs), and the MicrobeMod annotation pipeline.

